# Intuitive, reproducible high-throughput molecular dynamics in Galaxy: a tutorial

**DOI:** 10.1101/2020.05.08.084780

**Authors:** Simon A. Bray, Tharindu Senapathi, Christopher B. Barnett, Björn A. Grüning

## Abstract

This paper is a tutorial developed for the data analysis platform Galaxy. The purpose of Galaxy is to make high-throughput computational data analysis, such as molecular dynamics, a structured, reproducible and transparent process. In this tutorial we focus on 3 questions: How are protein-ligand systems parameterized for molecular dynamics simulation? What kind of analysis can be carried out on molecular trajectories? How can high-throughput MD be used to study multiple ligands? After finishing you will have learned about force-fields and MD parameterization, how to conduct MD simulation and analysis for a protein-ligand system, and understand how different molecular interactions contribute to the binding affinity of ligands to the Hsp90 protein.

## Introduction

Molecular dynamics (MD) is a commonly used method in computational chemistry and cheminformatics, in particular for studying the interactions between small molecules and large biological macromolecules such as proteins (1). However, the barrier to entry for MD simulation is high; not only is the theory difficult to master, but commonly used MD software is technically demanding. Furthermore, generating reliable, reproducible simulation data requires the user to maintain detailed records of all parameters and files used, which again poses a challenge to newcomers to the field. One solution to the latter problem is usage of a workflow management system such as Galaxy (2), which provides a selection of tools for molecular dynamics simulation and analysis (3). MD simulations are rarely performed singly; in recent years, the concept of high-throughput molecular dynamics (HTMD) has come to the fore (4, 5). Galaxy lends itself well to this kind of study, as we will demonstrate in this paper, thanks to features allowing construction of complex workflows, which can then be executed on multiple inputs in parallel.

This tutorial provides a detailed workflow for high-throughput molecular dynamics with Galaxy, using the N-terminal domain (NTD) of Hsp90 (heat shock protein 90) as a case-study. Galaxy (2) is a data analysis platform that provides access to thousands of tools for scientific computation. It features a web-based user interface while automatically and transparently managing underlying computation details. Galaxy makes high-throughput data analysis a structured, reproducible and transparent process. This tutorial provides sample data, workflows, hands-on material and references for further reading. It presumes that the user has a basic understanding of the Galaxy platform. The aim is to guide the user through the various steps of a molecular dynamics study, from accessing publicly available crystal structures, to performing MD simulation (leveraging the popular GROMACS (6, 7) engine), to analysis of the results.

The entire analysis described in this article can be conducted efficiently on any Galaxy server which has the needed tools. In particular, we recommend using the Galaxy Europe server (https://cheminformatics.usegalaxy.eu) or the Galaxy South Africa server (https://galaxy-compchem.ilifu.ac.za).

The tutorial presented in this article has been developed as part of the Galaxy Training Network (8) and its most up-to-date version is accessible online on the Galaxy Training Materials website (9), under the URL https://training.galaxyproject.org/training-material/topics/computational-chemistry/tutorials/htmd-analysis/tutorial.html.

### What is high-throughput molecular dynamics?

Molecular dynamics (MD) is a method to simulate molecular motion by iterative application of Newton’s laws of motion. It is often applied to large biomolecules such as proteins or nucleic acids. A common application is to assess the interaction between these macromolecules and a number of small molecules (e.g. potential drug candidates). This tutorial provides a guide to setting up and running a high-throughput workflow for screening multiple small molecules, using the open-source GROMACS tools provided through the Galaxy platform. Following simulation, the trajectory data is analyzed using a range of tools to investigate structural properties and correlations over time.

### Why is Hsp90 interesting to study?

The 90 kDa heat shock protein (Hsp90) is a chaperone protein responsible for catalyzing the conversion of a wide variety of proteins to a functional form; examples of the Hsp90 clientele, which totals several hundred proteins, include nuclear steroid hormone receptors and protein kinases (10). The mechanism by which Hsp90 acts varies between clients, as does the client binding site; the process is dependent on post-translational modifications of Hsp90 and the identity of co-chaperones which bind and regulate the conformational cycle (11).

Due to its vital biochemical role as a chaperone protein involved in facilitating the folding of many client proteins, Hsp90 is an attractive pharmaceutical target. In particular, as protein folding is a potential bottleneck to cellular reproduction and growth, blocking Hsp90 function using inhibitors which bind tightly to the ATP binding site of the NTD could assist in treating cancer; for example, the antibiotic geldanamycin and its analogs are under investigation as possible anti-tumor agents (12, 13).

In the structure which will be examined during this tutorial, the ligand of concern is a resorcinol, a common class of compounds with affinity for the Hsp90 N-terminal domain. It is registered in the PubChem database under the compound ID 135508238 (14). As can be seen by viewing the PDB structure, the resorcinol part of the structure is embedded in the binding site, bound by a hydrogen bond to residue aspartate-93. The ligand structure also contains a triazole and a fluorophenyl ring, which lie nearer to the surface of the protein.

## Methods: Simulation

In this section we will take the reader through the step-by-step process required to prepare, run and analyze Hsp90. A brief explanation of the theory and purpose of each step is provided. Refer to the hands-on sections as these describe the task with tools and parameters to be carried out using Galaxy.

### Get data

As a first step, we create a new Galaxy history and then we download a crystal structure for the Hsp90 protein from the Protein Data Bank (PDB). The structure is provided under accession code `6HHR` (16) and shows Hsp90 in complex with a ligand belonging to the resorcinol class.

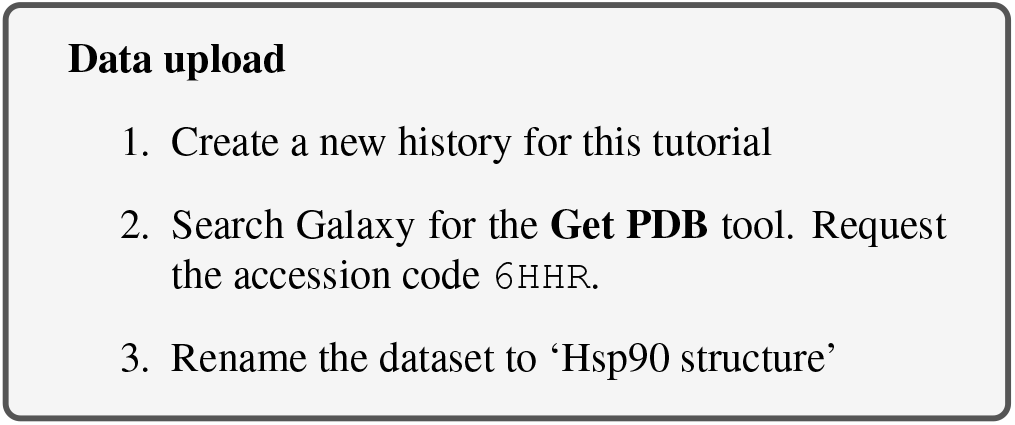

### Topology generation

Now we have downloaded a PDB structure of the protein we wish to study, we will start preparing it for MD simulation; this process may also be referred to as parameterization or topology generation.

GROMACS distinguishes between constant and dynamic attributes of the atoms in the system. The constant attributes (e.g. atom charges, bonds connecting atoms) are listed in the topology (TOP file), while dynamic attributes (attributes that can change during a simulation, e.g. atom position, velocities and forces) are stored in structure (PDB or GRO) and trajectory (XTC and TRR) files.

The PDB file we start from only explicitly states atom element (i.e. carbon, oxygen, and so on) and 3D Cartesian coordinates of each atom; additionally, it will usually not include hydrogen atoms. Therefore, before beginning simulation, we need to calculate the rest of the information contained within the topology file.

### Extract protein and ligand coordinates

Parameterization needs to be done separately for the ligand and protein. Therefore, the first step is to separate the PDB file into two sets of coordinates – one for the ligand and one for the protein. Here, we can make use of the simple text manipulation tools integrated into Galaxy.

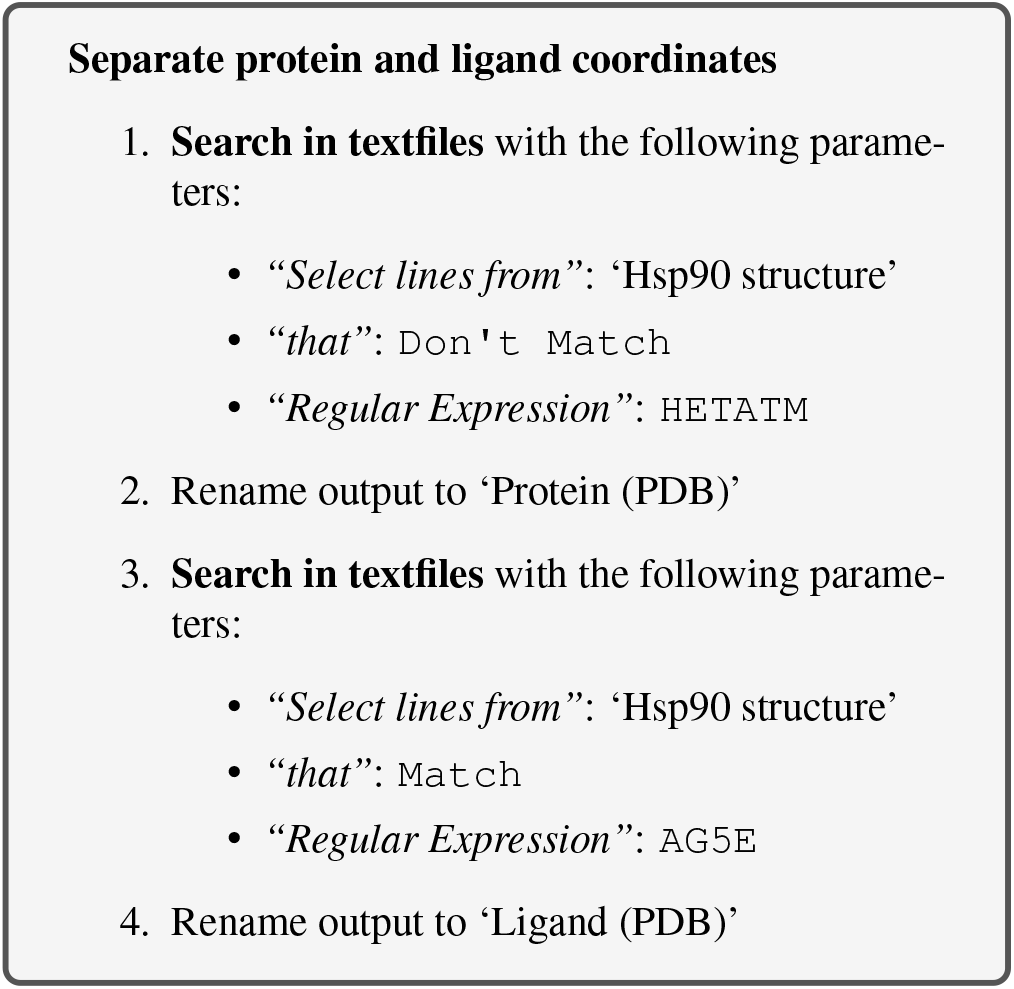

Here, we simply filter the original PDB twice: once for lines which do not match `HETATM`, which returns a PDB file containing only protein, not ligand and solvent; and once for lines which match the ligand’s identity code `AG5E`, which returns a PDB file containing only the ligand.

### Set up protein topology

Firstly, we need to calculate the topology for the protein file. We will use the **GROMACS initial setup** tool.

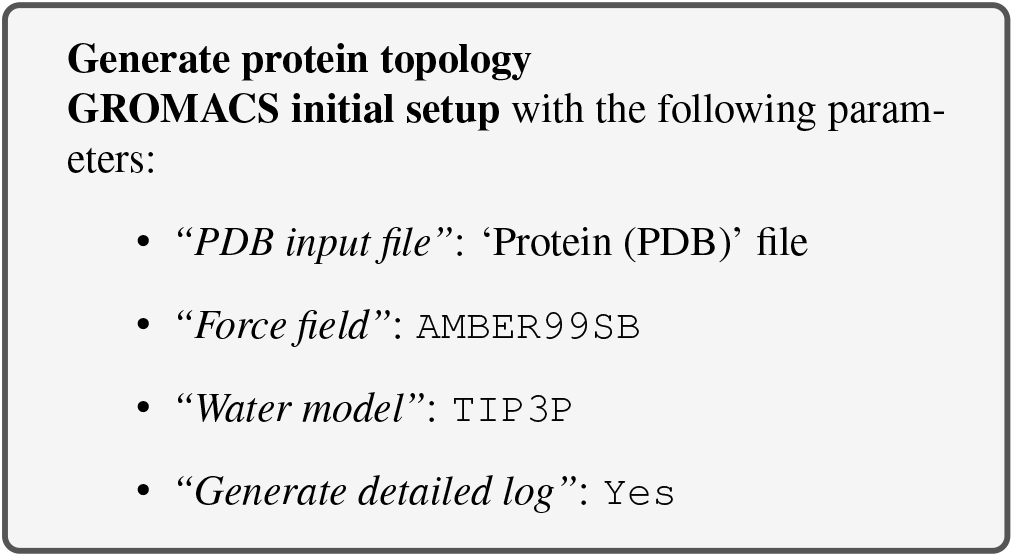

A force field is essentially a function to calculate the potential energy of a system, based on various empirical parameters (for the atoms, bonds, charges, dihedral angles and so on). There are a number of families of force fields; some of the most commonly used include CHARMM (17), AMBER (18), GROMOS (19) and OpenFF (20) (for a recent, accessible overview see (21)). We will use `ff99SB`, which is one of the main AMBER force fields for protein modeling.

Apart from the force field, a water model also needs to be selected to model the solvent; a wide range of models exist for this purpose. Here we are using the common TIP3P model, which is an example of a ‘three-site model’ – so-called because the molecule is modeled using three points, corresponding to the three atoms of water (Four-and five-site models include additional ‘dummy atoms’ representing the negative charges of the lone pairs of the oxygen atom) (22).

The tool produces four outputs: a GRO file (containing the coordinates of the protein), a TOP file (containing other information, including charges, masses, bonds and angles), an ITP file (which will be used to restrain the protein position in the equilibration step later on), and a log for the tool.

Please note all the GROMACS tools provided in Galaxy output a log. These provide useful information for debugging if we encounter any problems.

### Generate a topology for the ligand

To generate a topology for the ligand, we will use the **acpype**(23) tool. This provides a convenient interface to the AmberTools suite and allows us to easily create the ligand topology in the format required by GROMACS.

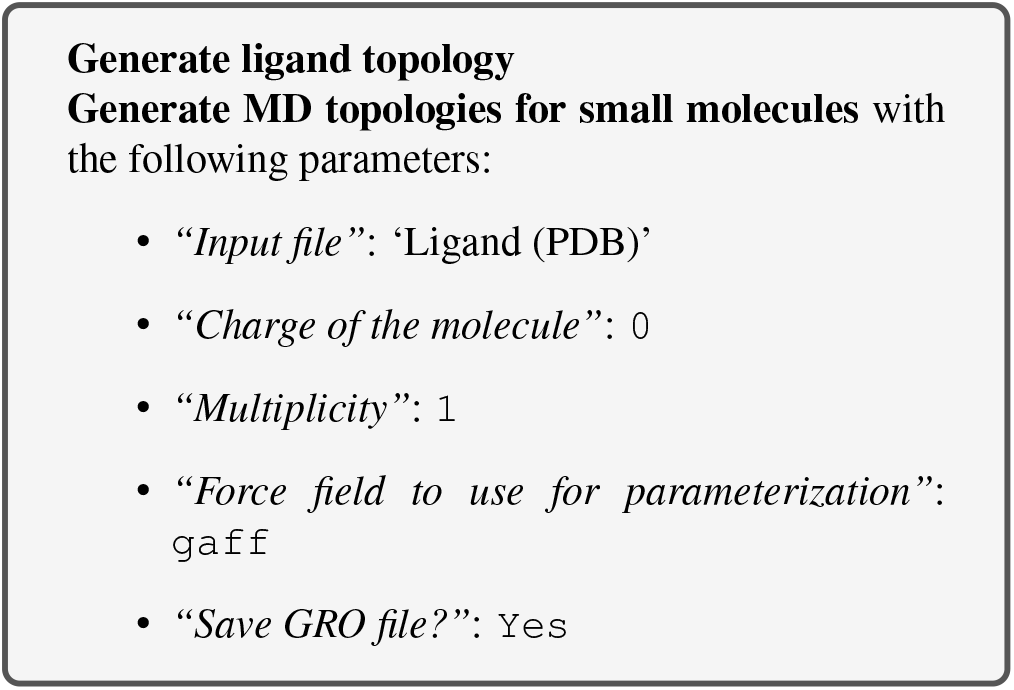

Here, we use GAFF (general AMBER force field), which is a generalized AMBER force field (24) which can be applied to almost any small organic molecule.

We select charge and multiplicity as appropriate. The ligand studied here is neutral, so the charge is 0. The multiplicity is 1, which will be the case for every simple organic molecule we encounter; only if we deal with more exotic species such as metal complexes or carbenes will we need to consider higher values.

Having generated topologies, we now need to combine them, define the box which contains the system, add solvent and ions, and perform an energy minimization step.

### Combine topology and GRO files

While we have separate topology and structure files for both protein and ligand, we need to combine them into a single set of files to continue with the simulation setup. A dedicated Galaxy tool is provided for this, using the Python library ParmEd (25).

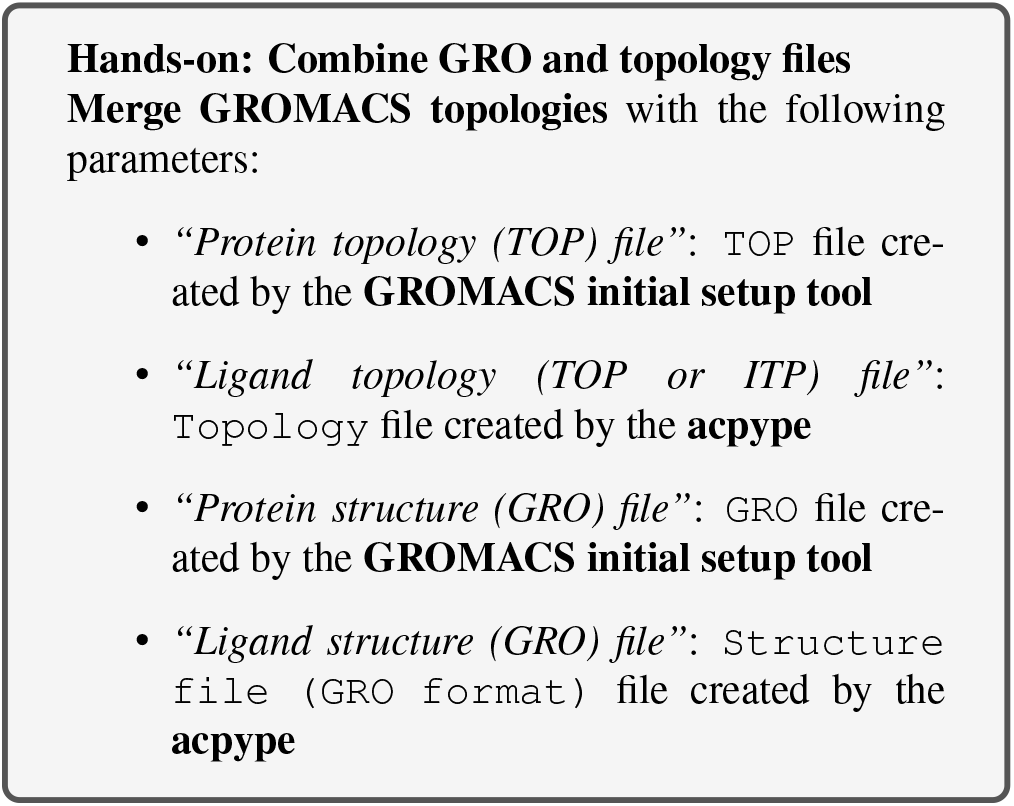

We now have a single GRO and TOP file, unifying the protein-ligand complex structure and topology.

### Create the simulation box with GROMACS structure configuration

The next step, once combined coordinate (GRO) and topology (TOP) files have been created, is to create a simulation box in which the system is situated.

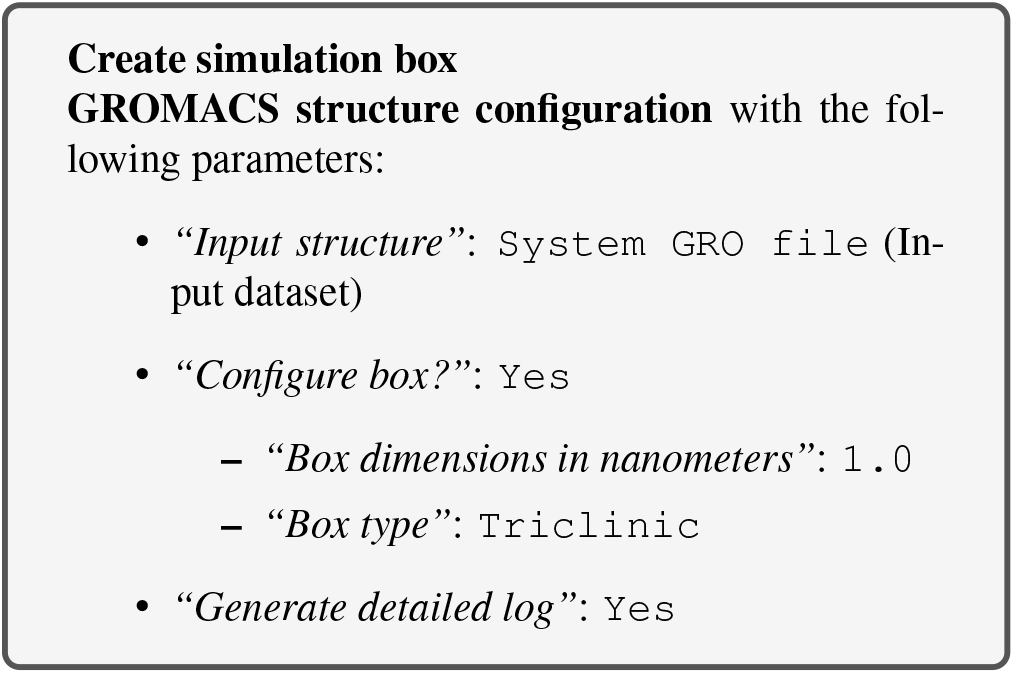

This tool returns a new GRO structure file, containing the same coordinates as before, but defining a simulation box such that every atom is a minimum of 1 nm from the box boundary. A variety of box shapes are available to choose: we select triclinic, as it provides the most efficient packing in space and thus fewer computational resources need to be devoted to simulation of solvent.

### Solvation

The next step is solvation of the newly created simulation box – as we are simulating under biological conditions, we use water as the solvent. Note that the system is charged (depending on the pH) – the solvation tool also adds sodium or chloride ions (replacing existing water molecules) as required to neutralize this.

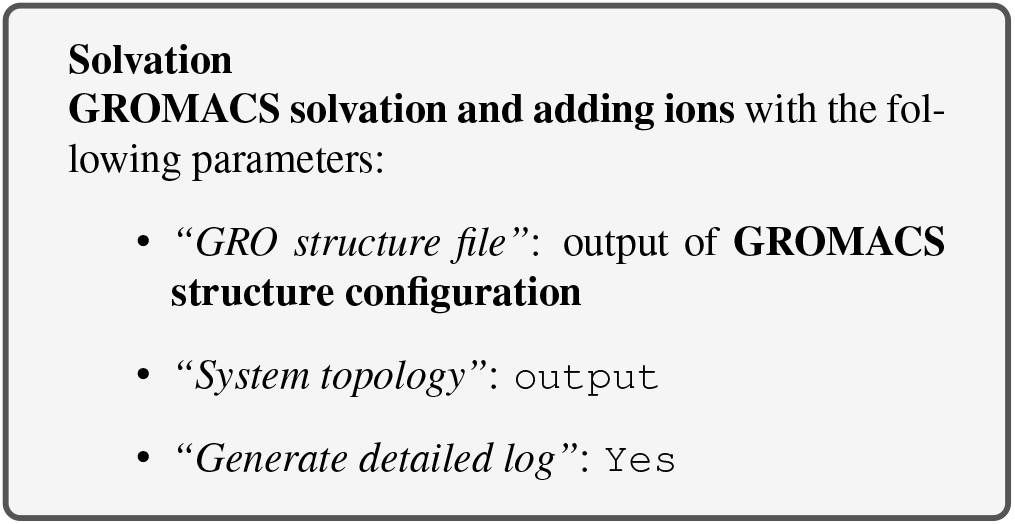

### Energy minimization

After the solvation step, parameterization of the system is complete and preparatory simulations can be performed. The first of theses is energy minimization, which can be carried out using the **GROMACS energy minimization** tool. The purpose of energy minimization is to relax the structure, removing any steric clashes or unusual geometry which would artificially raise the energy of the system.

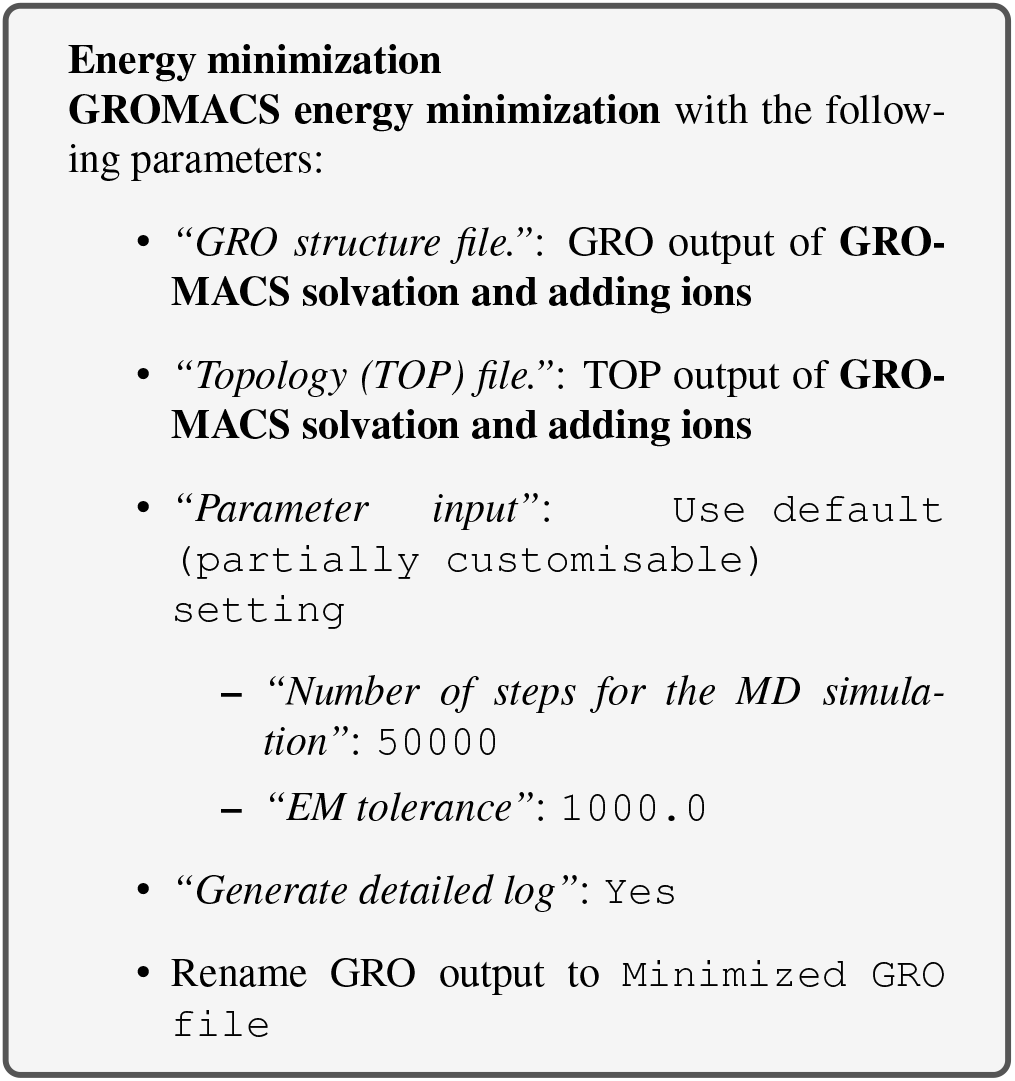

The EM tolerance here refers to the maximum force which will be allowed in a minimized system. The simulation will be terminated when the maximum force is less than this value, or when 50000 steps have elapsed.

As an aside, we can use the ‘Extract energy components’ tool to plot the convergence of the potential energy during the minimization.

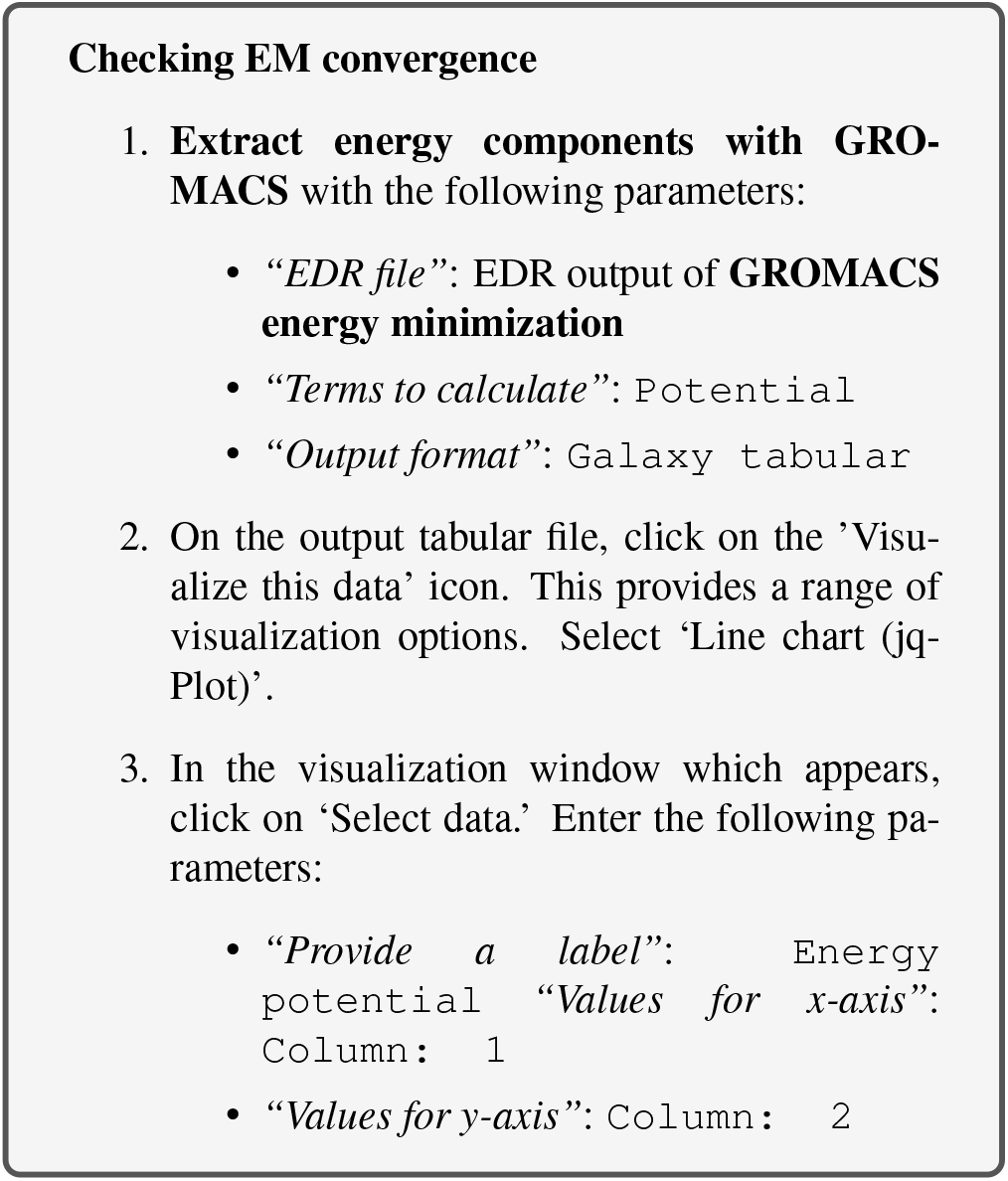

The resulting plot should resemble Figure 2. The system first drops rapidly in energy, before slowly converging on the minimized state.

**Fig. 1.**
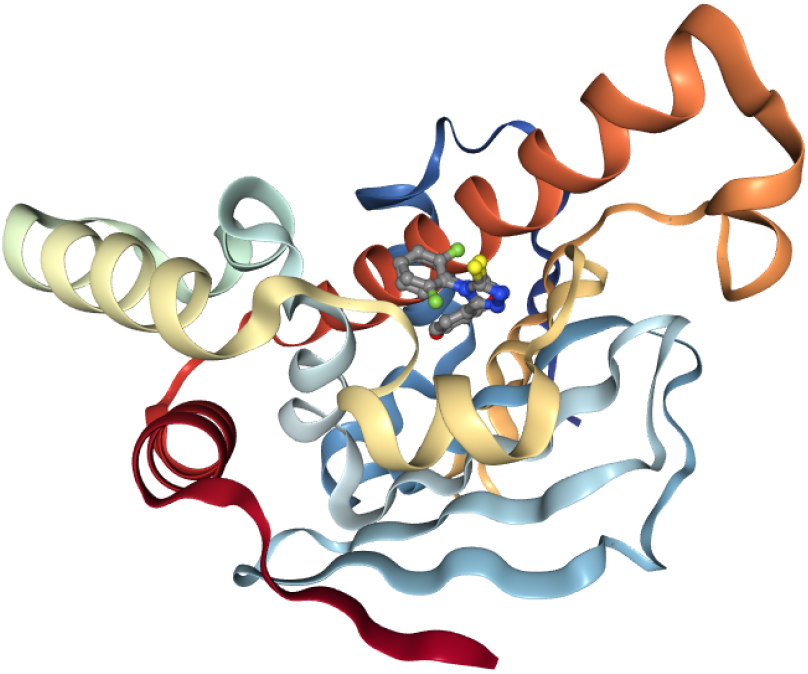
Hsp90 cartoon view. Hsp90 cartoon with ligands in active site, rendered using the Galaxy NGL plugin (15)

**Fig. 2.**
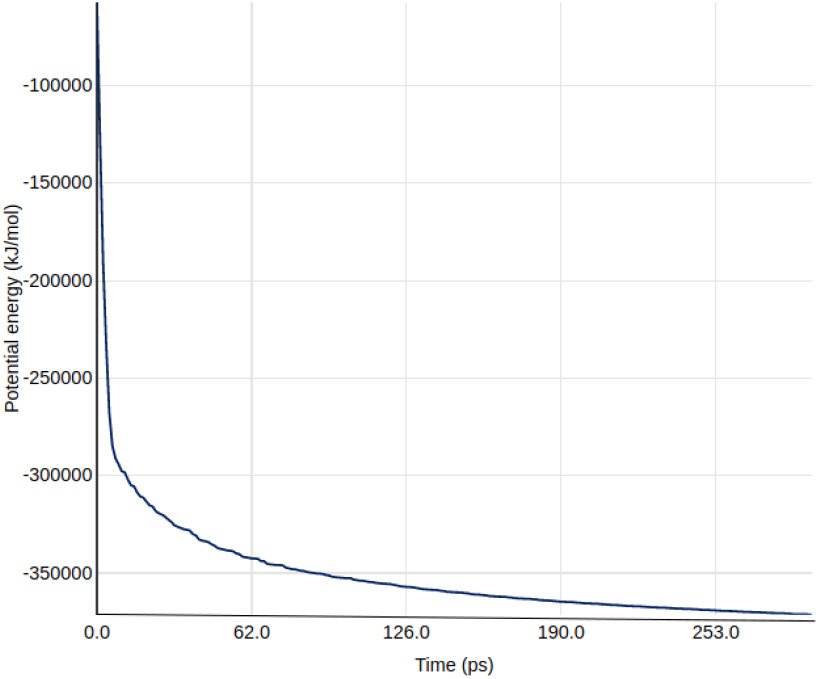
Energy potential during the EM simulation.

### Equilibration

At this point equilibration of the solvent around the solute (i.e. the protein) is necessary. This is performed in two stages: equilibration under an NVT (or isothermal-isochoric) ensemble, followed by an NPT (or isothermal-isobaric) ensemble. Use of the NVT ensemble entails maintaining constant number of particles, volume and temperature, while the NPT ensemble maintains constant number of particles, pressure and temperature.

For equilibration, the protein must be held in place while the solvent is allowed to move freely around it. This is achieved using the position restraint file (ITP) we created in system setup. When we specify this restraint, protein movement is not forbidden, but is energetically penalized.

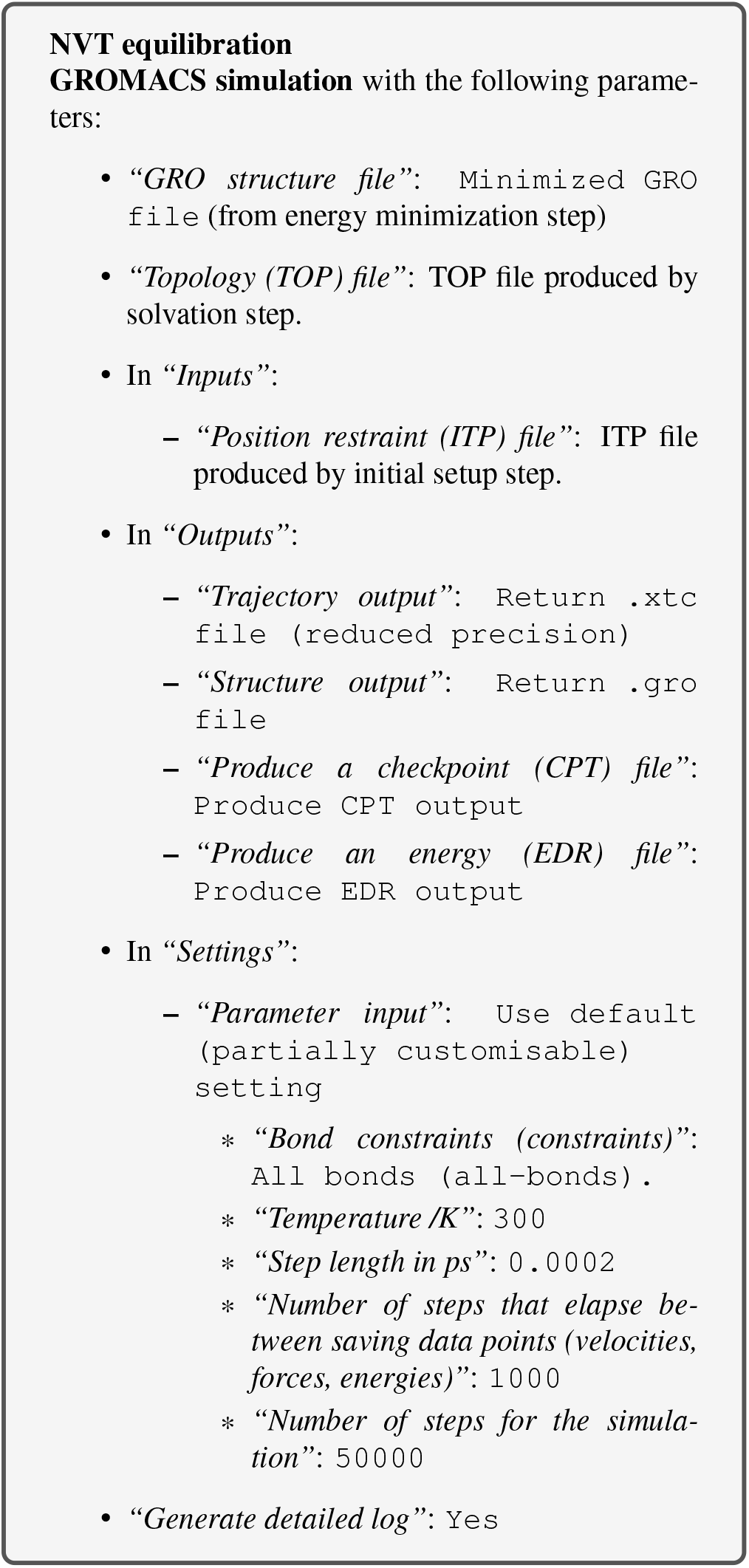

Once the NVT equilibration is complete, it is worth using the **Extract energy components** tool again to check whether the system temperature has converged on 300 K. This can be done as described for energy minimization, this time specifying `Temperature` under *Terms to calculate* rather `Potential`. The plot should show the temperature reaching 300 K and remaining there, albeit with some fluctuation.

Having stabilized the temperature of the system with NVT equilibration, we also need to stabilize the pressure of the system. We therefore equilibrate again using the NPT (constant number of particles, pressure, temperature) ensemble.

Note that we can continue where the last simulation left off (with new parameters) by using the checkpoint (CPT) file saved at the end of the NVT simulation.

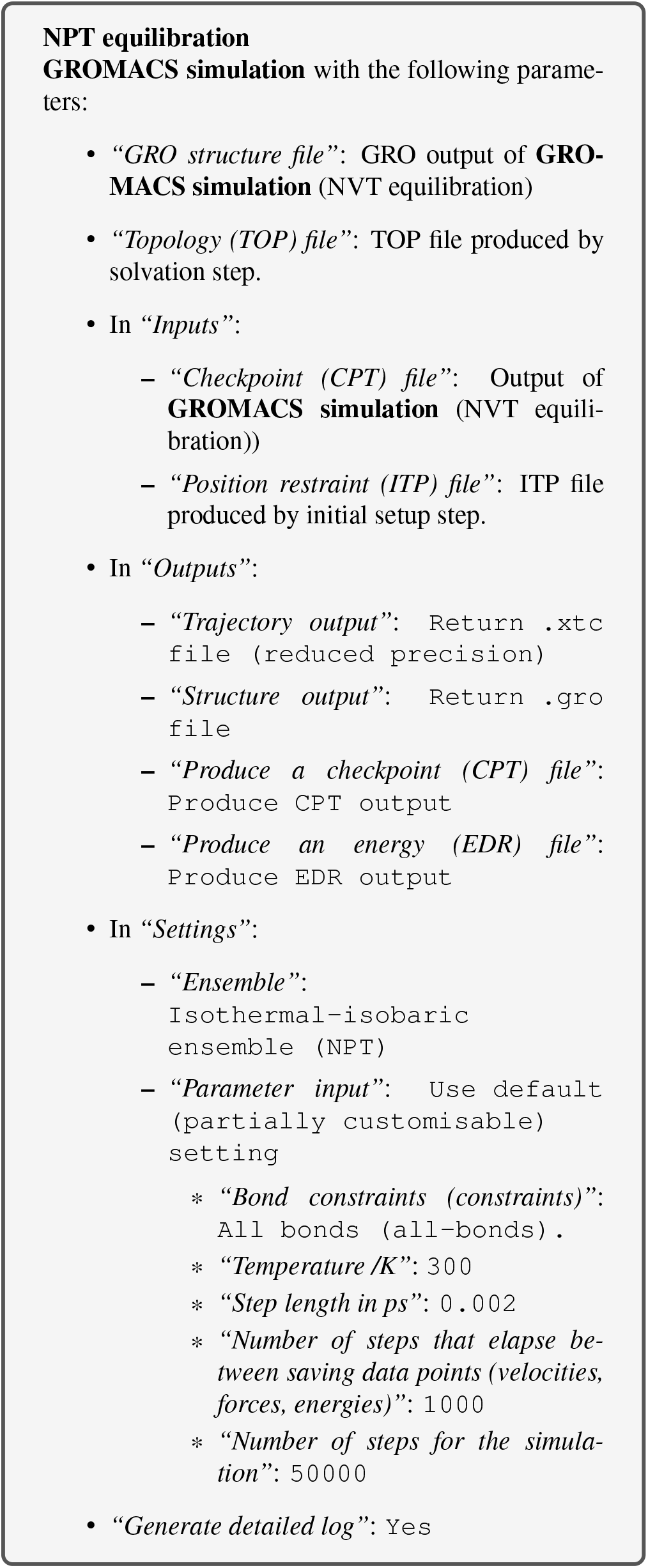

After the NPT equilibration is complete, **Extract energy components** can be used again to view the pressure of the system. This is done as described for energy minimization, specifying `Pressure` under *Terms to calculate*. The plot should show convergence on 1 bar and remaining there, although some fluctuation is expected.

### Production simulation

We can now remove the restraints and continue with the production simulation. The simulation will run for 1 million steps, with a step size of 1 fs, so will have a total length of 1 ns. This is rather short compared to the state-of-the-art, but sufficient for the purposes of a tutorial. For longer-scale simulations, the tool can be used multiple times (with the checkpoint file) to continue the existing simulation.

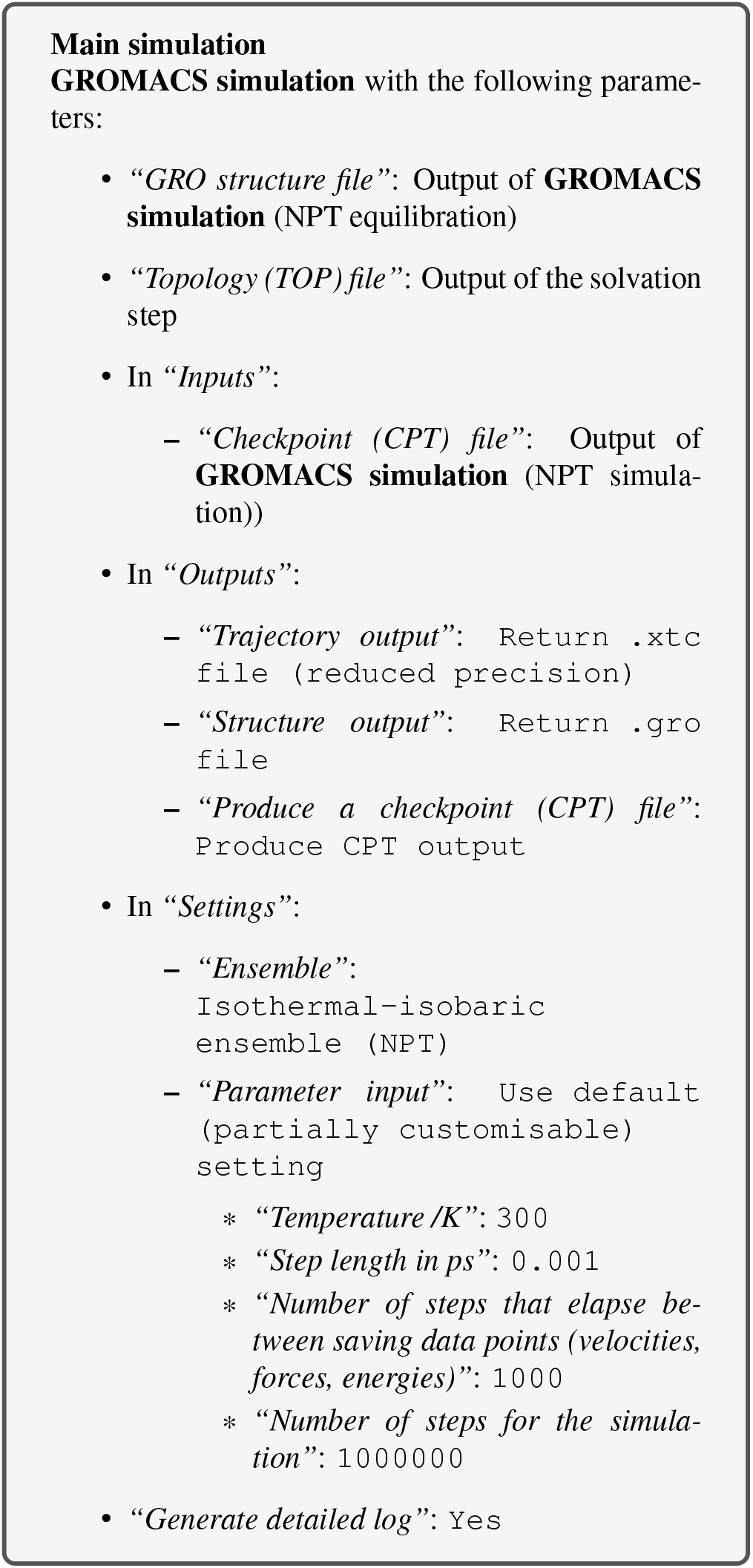

## Methods: Analysis

An analysis of the GROMACS simulation outputs (structure and trajectory file) will be carried out using Galaxy tools developed for computational chemistry (3) based on popular analysis software, such as MDAnalysis (26), MDTraj (27), and Bio3D (28). These tools output both tabular files as well as a variety of attractive plots.

### Convert coordinate and trajectory formats

Before beginning a detailed analysis, the structure and trajectory files generated previously need to be converted into different formats. First, convert the structural coordinates of the system in GRO format into PDB format. This PDB file will be used by most analysis tools as a starting structure. This tool can also be used to carry out initial setup (as discussed in the simulation methods section) for GROMACS simulations and convert from PDB to GRO format. Next, convert the trajectory from XTC to DCD format, as a number of tools (particularly those based on Bio3D) only accept trajectories in DCD format. This tool can also be used to interconvert between several other trajectory formats.

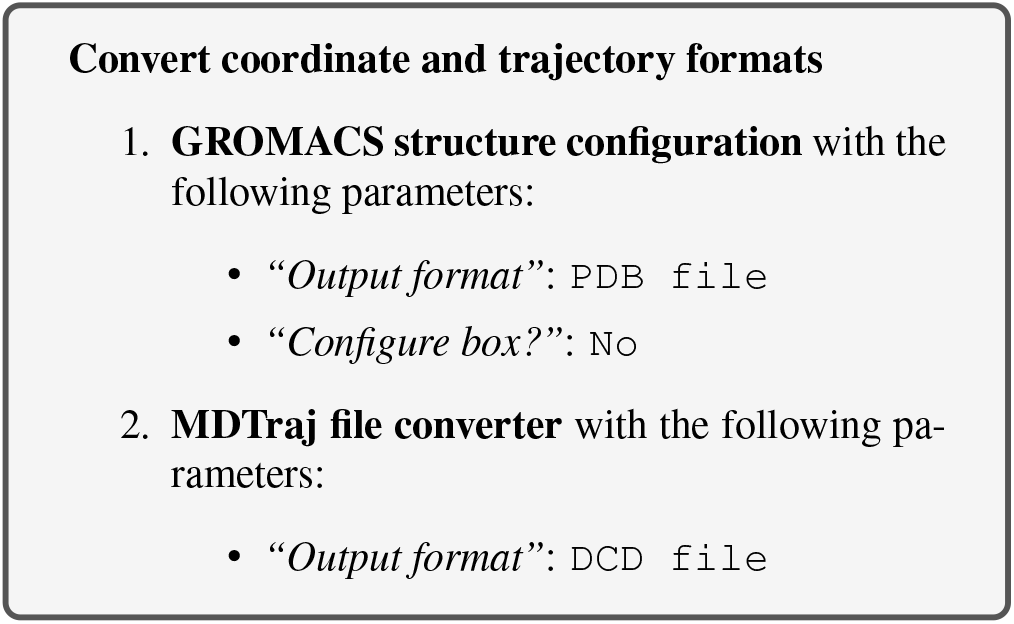

### RMSD analysis

The Root Mean Square Deviation (RMSD) and Root Mean Square Fluctuation (RMSF) are calculated to check the stability and conformation of the protein and ligand through the course of the simulation. RMSD is a standard measure of structural distance between coordinate sets that measures the average distance between a group of atoms. The RMSD of the C*α* atoms of the protein backbone is calculated here and is a measure of how much the protein conformation has changed between different time points in the trajectory. Note that for more complex systems, you may need to consider a more focused selection.

For the RMSD analysis of the ligand, the ‘Select domains’ parameter of the tool can for convenience be set to ‘Ligand’; however, this automatic selection sometimes fails. The other alternative, which we apply here, is to specify the ‘Residue ID’ in the textbox provided. In this example the ligand’s Residue ID is ‘G5E’. The output is the requested RMSD data as a time series, the RMSD plotted as a time series and as a histogram (for example, see Figure 3 in the results section).

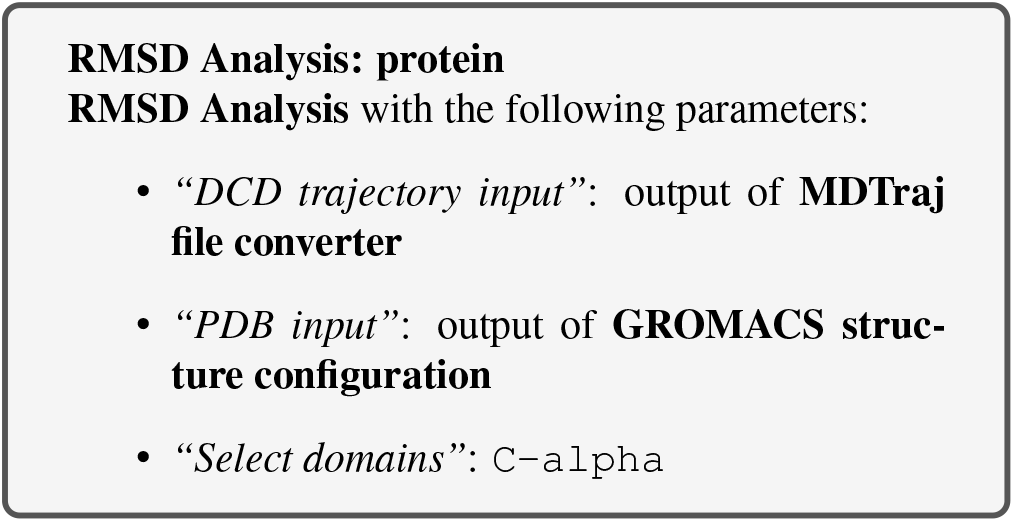

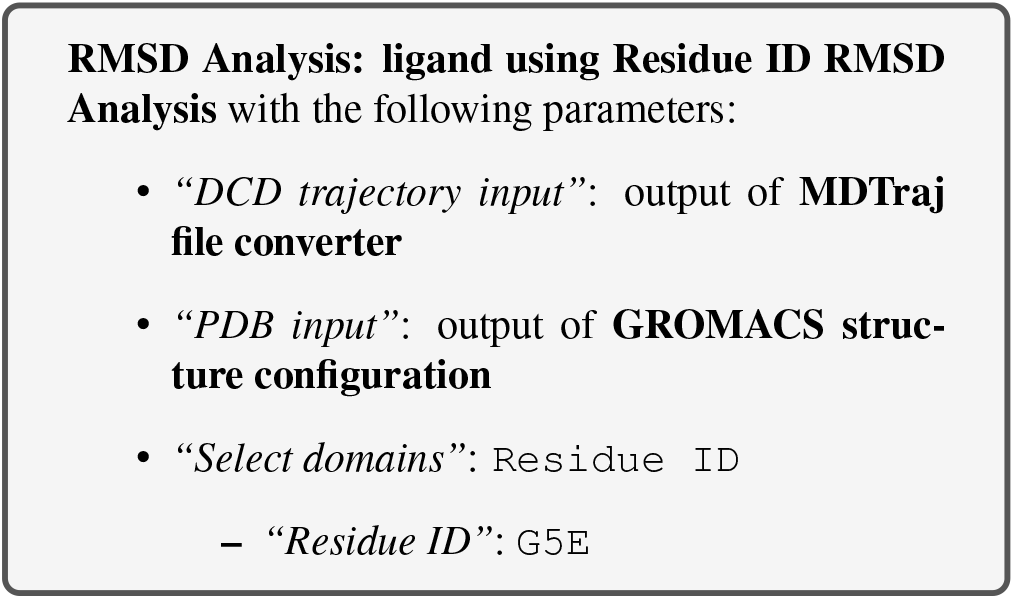

**Fig. 3.**
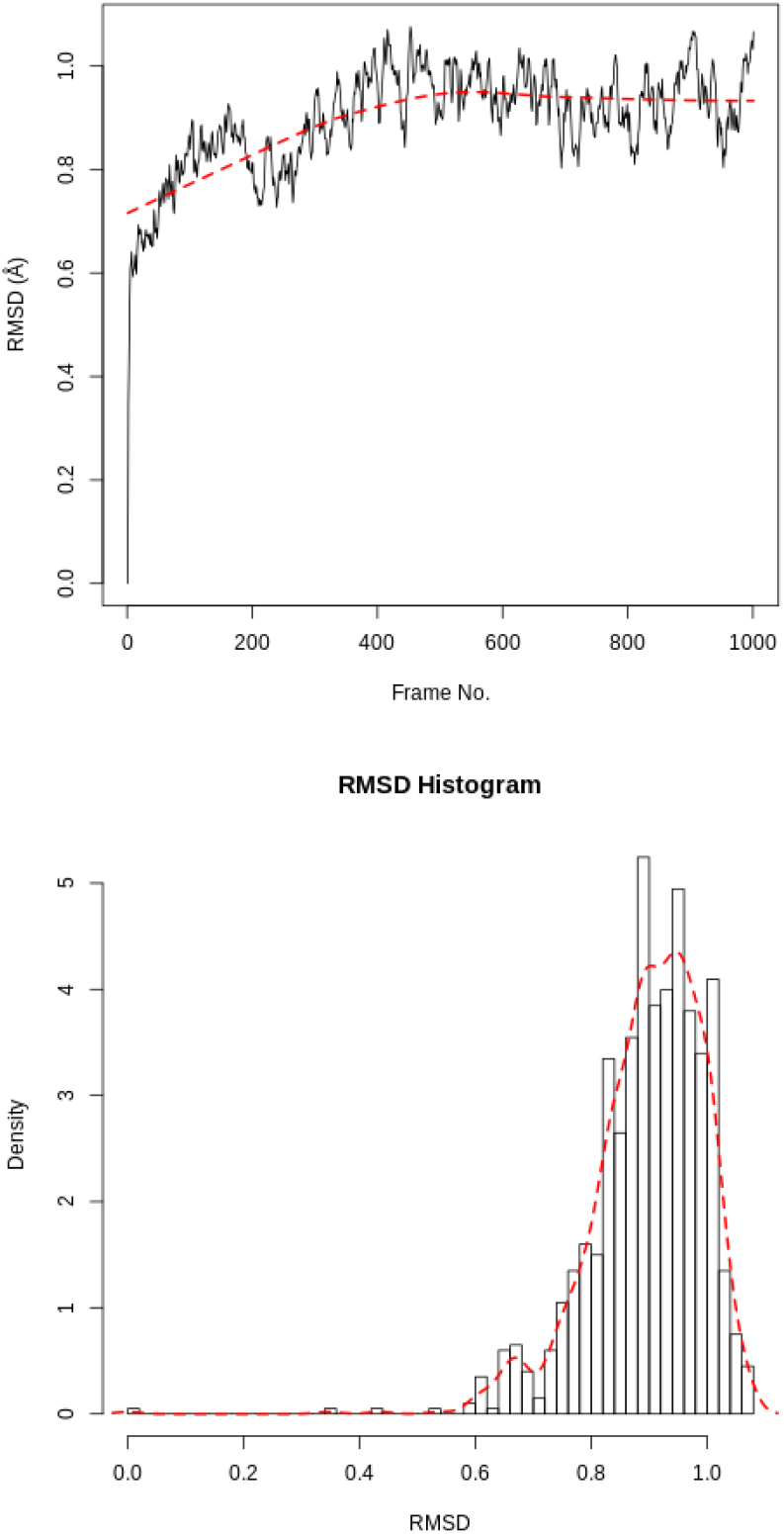
RMSD for protein. RMSD time series and histogram for the protein.

### RMSF analysis

The Root Mean Square Fluctuation (RMSF) is valuable to consider, as it represents the deviation at a reference position over time. The fluctuation in space of particular amino acids in the protein are considered. The C*α* of the protein, designated by `C-alpha`, is a good selection to understand the change in protein structure. Depending on the system these fluctuations can be correlated to experimental techniques including Nuclear Magnetic Resonance (NMR) and Mössbauer spectroscopy (29, 30). The output from the tools is the requested RMSF data and the RMSF plotted as a time series (for example, see Figure 5 in the results section).

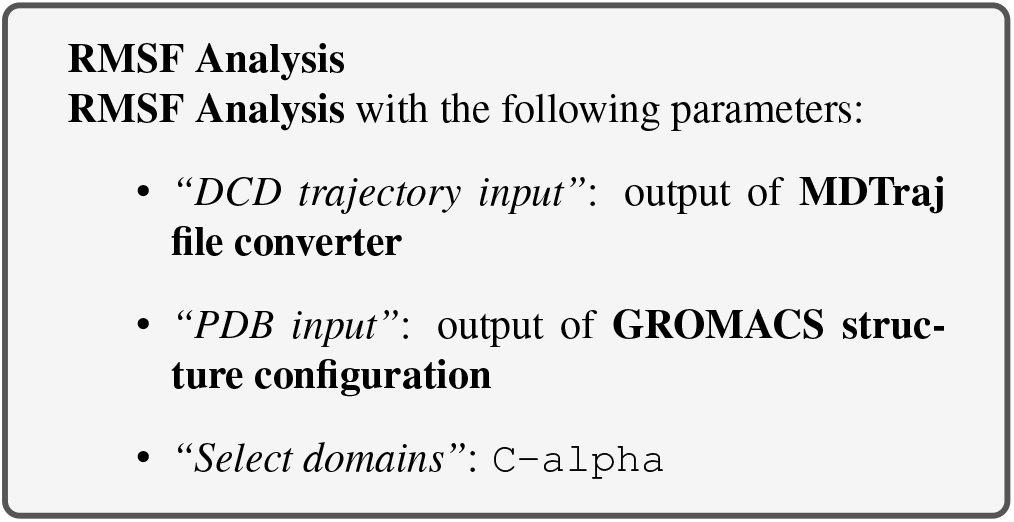

### PCA

Principal component analysis (PCA) converts a set of correlated observations (movement of selected atoms in protein) to a set of principal components (PCs) which are linearly independent (or uncorrelated). Here several related tools are used. The PCA tool calculates the PCA in order to determine the relationship between statistically meaningful conformations (major global motions) sampled during the trajectory. The C*α* carbons of the protein backbone are again a good selection for this purpose. Outputs include the PCA raw data and figures of the relevant principal components (PCs) as well as an eigenvalue rank plot (see Figure 6) which is used to visualize the proportion of variance due to each principal component (remembering that the PCs are ranked eigenvectors based on the variance). Having discovered the principal components usually these are visualized. The PCA visualization tool will create trajectories of specific principal components which can be viewed in a molecular viewer such as VMD (31) or NGL viewer (15). We also consider the PCA cosine content which when close to 1 indicates that the simulation is not converged and a longer simulation is needed. For values below 0.7, no statement can be made about convergence or lack thereof.

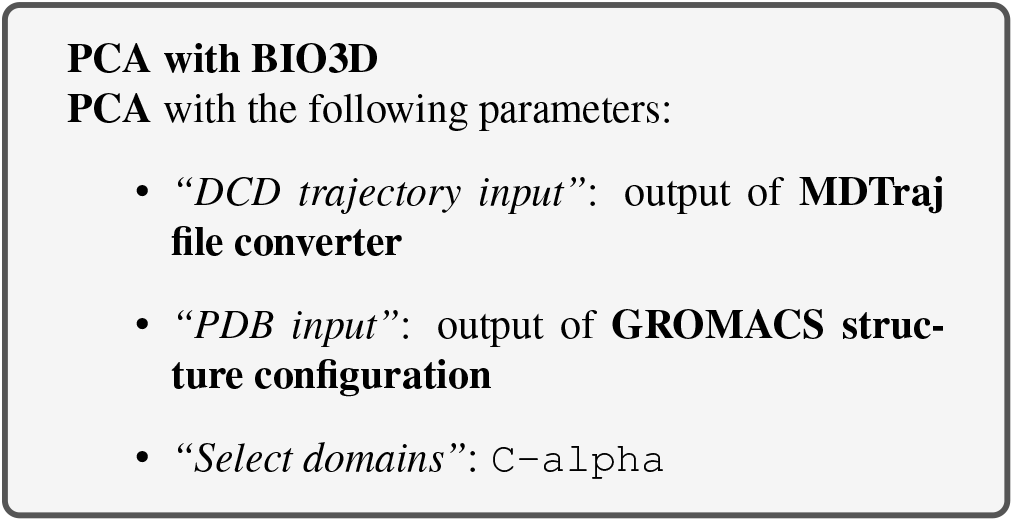

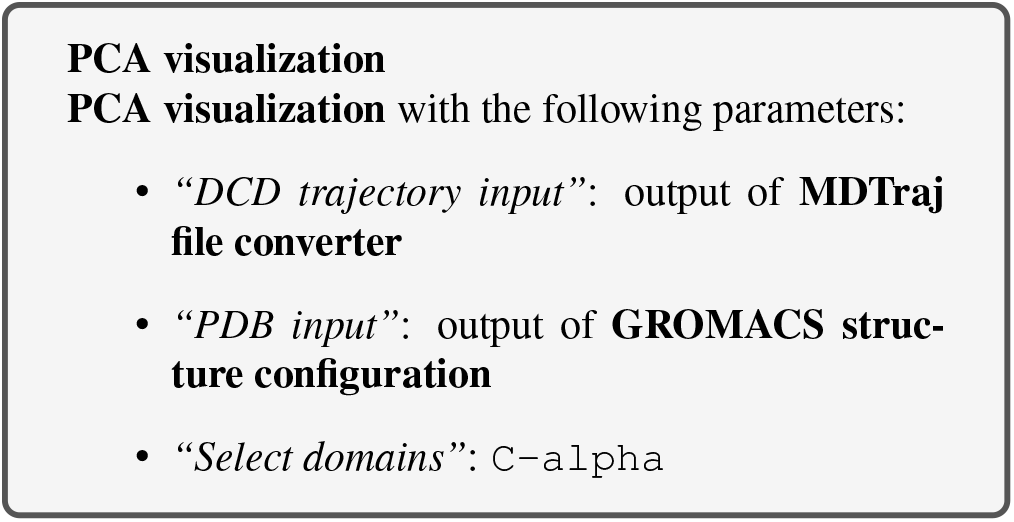

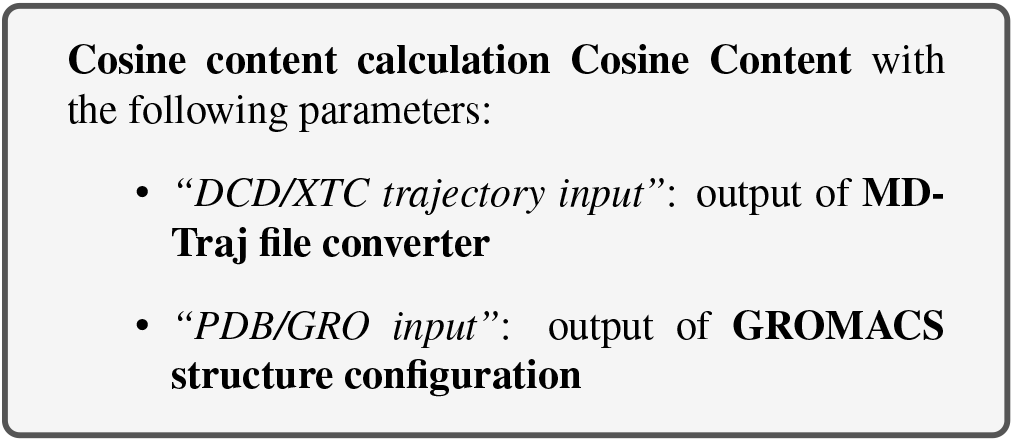

### Hydrogen bond analysis

Hydrogen bonding interactions contribute to binding and are worth investigating, in particular persistent hydrogen bonds. All possible hydrogen bonding interactions between the two selected regions, here the protein and the ligand, are investigated over time using the VMD hydrogen bond analysis tool included in Galaxy. Hydrogen bonds are identified and in the output the total number of hydrogen bonds and occupancy over time is returned.

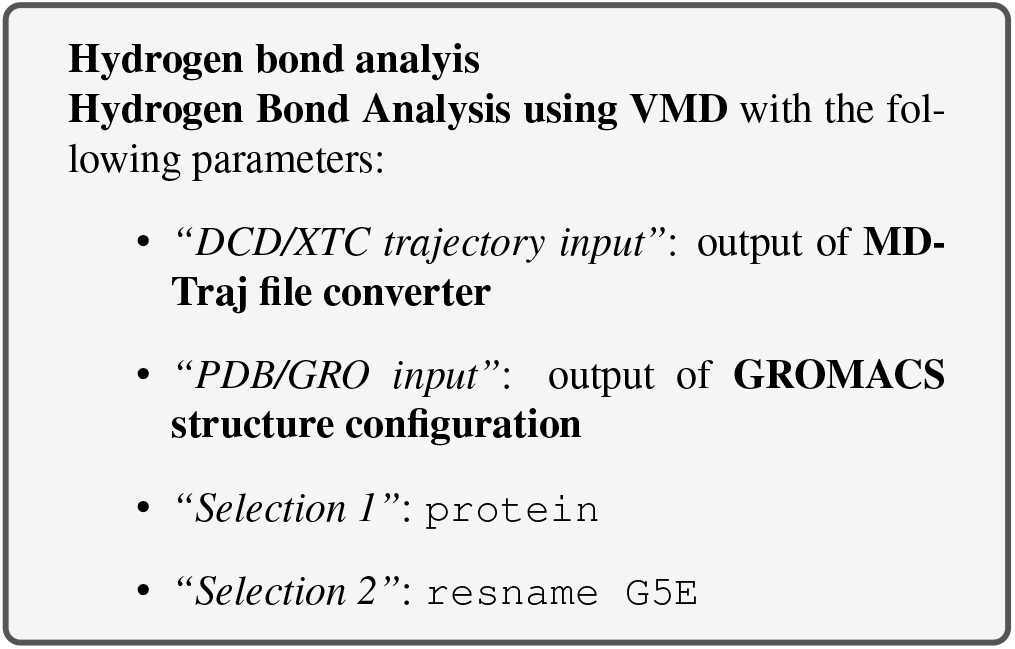

## Results and Discussion

After the completion of the simulation, the following questions arise: 1) is the simulation converged enough, and 2) what interesting molecular properties are observed. The timescale of motions of interest are in the picosecond to nanosecond range; these are motions such as domain vibration, hydrogen bond breaking, translation diffusion and side chain fluctuations. To observe meaningful conformational transitions of the protein *μ*s sampling would be needed, but this is not the purpose here.

The PCA cosine content of the dominant motion related to PC1 is 0.93, indicating that the simulation is not fully converged. This is expected due to the short simulation length. For production level simulations, it is the norm to extend simulations to hundreds of nanoseconds in length, if not microseconds. As this tutorial is designed to be carried out on public webservers, we limit simulations to 1 ns, as we cannot provide a large amount of computational resources for training purposes.

### RMSD protein

The RMSD time series for the protein shows a thermally stable and equilibrated structure that plateaus at 1.0Å with an average RMSD between 0.8Å and 1.0Å. There are no large conformational changes during the simulation. The RMSD histogram confirms this, see Figure 3. Note these graphs are automatically created by Galaxy as part of the tool’s outputs.

### RMSD ligand

Calculating the RMSD of the ligand is necessary to check if it is stable in the active site and to identify possible binding modes. If the ligand is not stable, there will be large fluctuations in the RMSD.

In our case the ligand is stable with a single binding mode. The RMSD fluctuates around 0.3Å, with a slight fluctuation near the end of the simulation. This is more clearly seen in the histogram, see Figure 4. The conformation seen during simulation is very similar to that in the crystal structure and the ligand is stable in the active site.

**Fig. 4.**
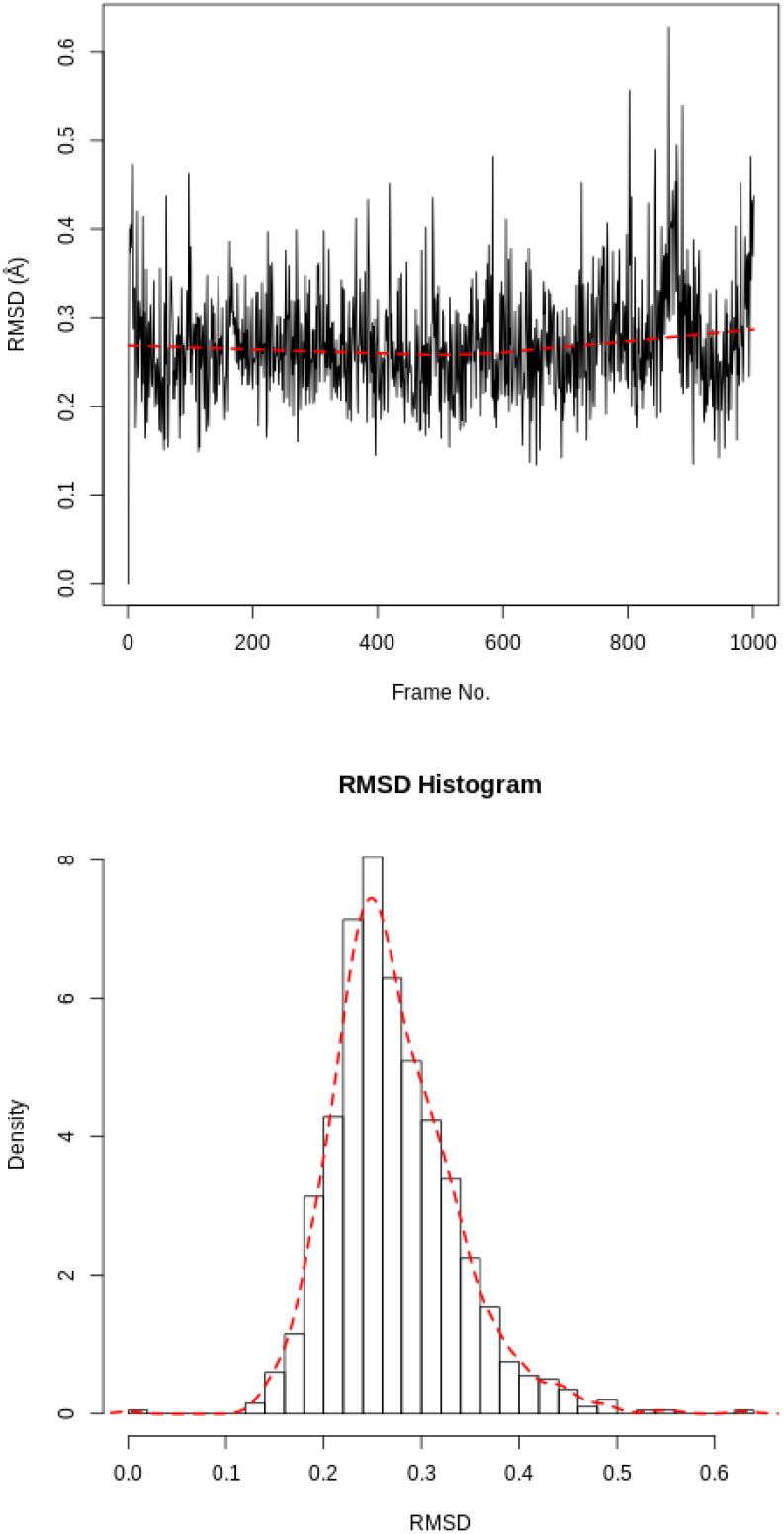
RMSD for the ligand. RMSD time series and histogram for the ligand.

### RMSF

When considering the RMSF (Figure 5), fluctuations greater than 1.0Å are of interest; for example see the fluctuations near residue positions 50, 110 and 160. Inspecting the structure with molecular visualization software such as VMD, these can be seen to correspond to flexible loop regions on the protein surface. In addition, very large fluctuations are seen for the C-terminus; this is common and no investigation is needed.

**Fig. 5.**
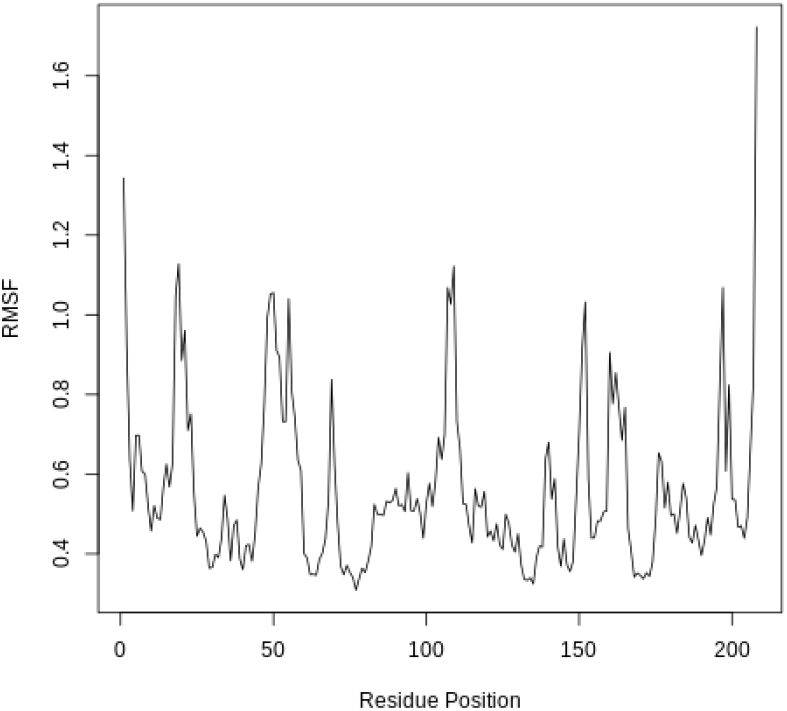
RMSF for the protein. RMSF(Å) vs the residue position. Large fluctuations occur at various positions, which correspond to flexible loop regions on the surface of the protein.

Note that the first few residues of this protein are missing in the PDB, and therefore residue position 0 in the RMSF corresponds to position 17 in the Hsp90 FASTA primary sequence. This is a fairly common problem that can occur with molecular modeling of proteins, where there may be missing residues at the beginning or within the sequence.

### PCA

The first three principal components are responsible for 32.8% of the total variance, as seen in the eigenvalue rank plot (Figure 6). The first principal component (PC1) accounts for 15.4% of the variance (see PC1 vs PC2 and eigenvalue rank plots in Figure 6). Visualization of PC1 using VMD shows a rocking motion and wagging of the C-terminus.

**Fig. 6.**
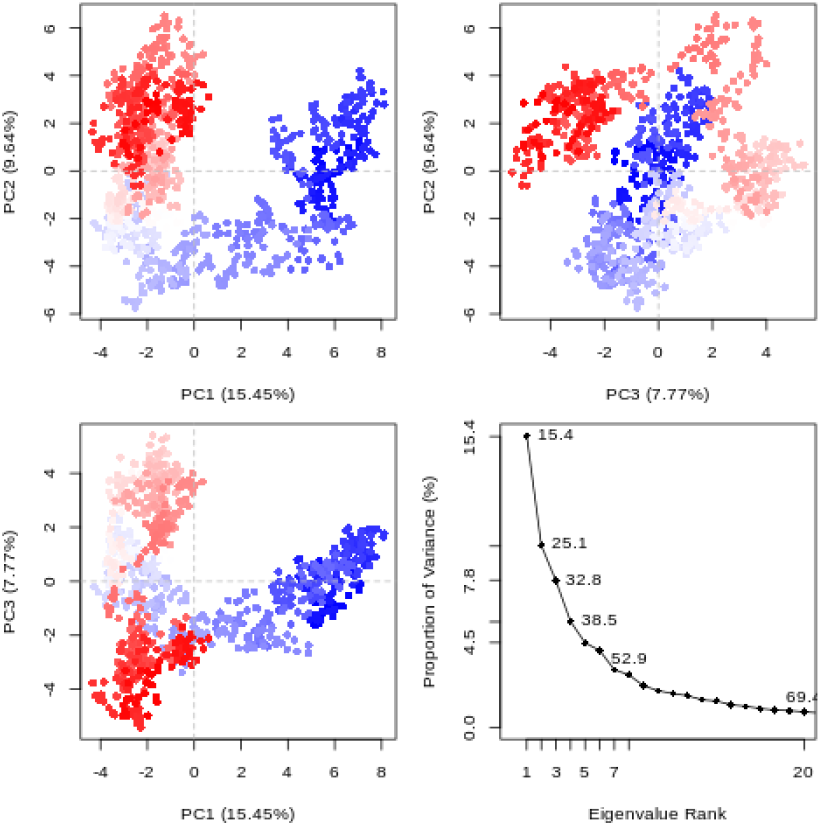
Principal Component Analysis. PCA results which include graphs of PC2 vs PC1, PC2 vs PC3, PC3 vs PC1 colored from blue to red in order of time, and an eigenvalue rank plot. In the eigenvalue plot the cumulative variance is labeled for each data point.

### Hydrogen bonding

The active site of this protein is quite hydrophobic, yet multiple hydrogen bonds were identified. The hydrogen bond between aspartate-93 and the ligand (as identified in the crystal structure) was found to be persistent, meeting the hydrogen bond criteria for 89.22% of the simulation. A hydrogen bond between the ligand and the carbonyl group of glycine-97 was found to have a 15.27% occupancy. Hydrogen bonding interactions with threonine-184, asparagine-51 and lysine-58 were also observed but these are not persistent and only present for a minority of the simulation. These values can be accessed from the ‘Percentage occupancy of the H-bond’ output of the hydrogen bond analysis tool.

## High throughput workflows

Up until this step, Galaxy tools have been applied sequentially to datasets. This is useful to gain an understanding of the steps involved, but becomes tedious if the workflow needs to be run on multiple protein-ligand systems. Fortunately, Galaxy allows entire workflows to be executed with a single mouse-click, enabling straightforward high-throughput analyses.

We will demonstrate the high-throughput capabilities of Galaxy by running the workflow detailed so far on a further three ligands.

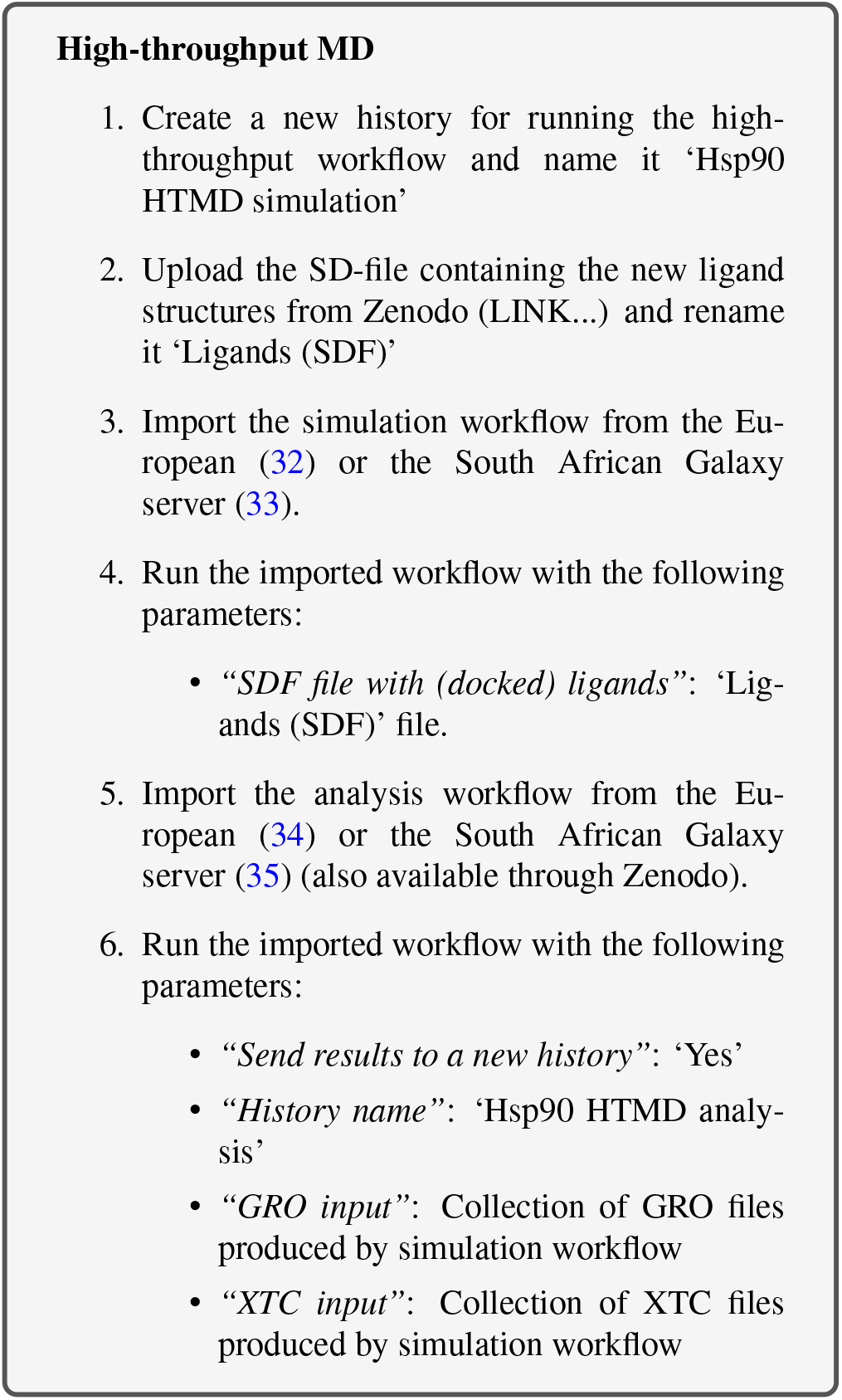

This process runs the entire simulation and analysis procedure described so far on the new set of ligands. It uses Galaxy’s collection (36) feature to organize the data; each item in the history is a collection (essentially a directory containing multiple individual datasets) containing one file corresponding to each of the input ligands.

Note that the SD-file needs to contain ligands with the correct 3D coordinates for MD simulation. The easiest way to obtain these is using a molecular docking tool such as Autodock Vina (37) or rDock (38); tutorials and workflows are available for both of these from the Galaxy Training Network. As an example, the history in which the SD-file used in the HTMD workflow is generated (using AutoDock Vina) is provided (39).

### Further information

Apart from manual setups or collections, there are several other alternatives which are helpful in scaling up workflows. Galaxy supports and provides training material for converting histories to workflows (40), using multiple histories (41), and the Galaxy Application Programming Interface (API) (42). For beginners and users who prefer a visual interface, automation can be done using multiple histories and collections with the standard Galaxy user interface.

If you are able to write small scripts, you can automate everything you have learned here with the Galaxy API. This allows you to interact with the server to automate repetitive tasks and create more complex workflows (which may have repetition or branching). The simplest way to access the API is through the Python library BioBlend (43). An example Python script, which uses BioBlend to run the GROMACS simulation workflow for each of a list of ligands, is given in the hands-on box below.

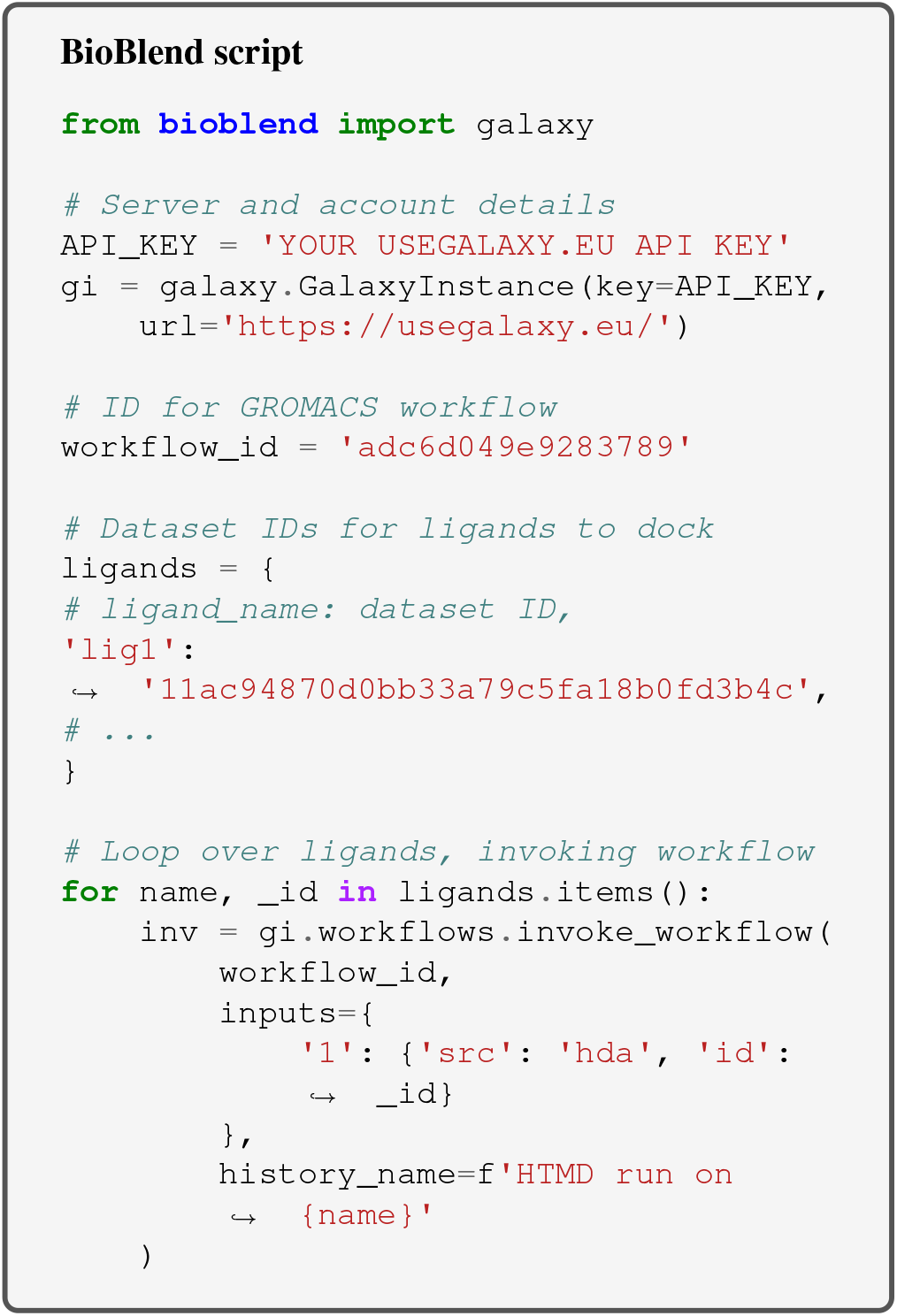

## Conclusion

This tutorial provides a guide on how to study protein-ligand interaction using molecular dynamics in Galaxy. Performing such analyses in Galaxy makes it straightforward to set up, schedule and run workflows, removing much of the difficulty from MD simulation. Thus, the technical barrier to performing high-throughput studies is greatly reduced. Results are structured in the form of Galaxy histories or collections, and include ready-plotted diagrams, which ensure data can be easily understood and reproduced if necessary. Apart from streamlining the process for existing MD users, this tutorial should also prove useful as a pedagogical guide for educating students or newcomers to the field.

After completing the tutorial, the user will be familiar at a basic level with a range of MD analysis techniques, and understand the steps required for a typical MD simulation. Thus, they will be equipped to apply these tools to their own problems.

### Funding

This work was supported by funding from the following organizations: S.A.B. was funded by the European Open Science Cloud (EOSC-Life) (Grant No. 824087); T.S. and C.B.B. were funded by the University of Cape Town’s Research Committee (URC) and by the National Research Foundation of South Africa (Grant Numbers 115215 and 116362); and was funded by the German Research Foundation for the Collaborative Research Center 992 Medical Epigenetics [SFB 992/1 2012 and SFB 992/2 2016].

### Data and material availability

Data and materials are available on GitHub:

- European Galaxy server (https://cheminformatics.usegalaxy.eu)
- Galaxy Computational Chemistry South Africa server (https://galaxy-compchem.ilifu.ac.za)
- Galaxy Training Network website (https://training.galaxyproject.org/topics/computational-chemistry/tutorials/htmd-analysis/tutorial.html)
- Supplementary Material, including workflows and data used (https://github.com/galaxycomputationalchemistry/htmd-paper-sm)

## Acknowledgements

We thank Bérénice Batut for helpful comments and discussions. In addition, we thank the entire Galaxy Training Network, as well as the European Galaxy and ilifu teams, for their support.

The European Galaxy server, which was used for the calculations described, is in part funded by Collaborative Research Centre 992 Medical Epigenetics (DFG grant SFB 992/1 2012) and German Federal Ministry of Education and Research (BMBF grants 031 A538A/A538C RBC, 031L0101B/031L0101C de.NBI-epi (de.NBI)).

